# Gαq Mediates Clozapine Effects in *Caenorhabditis* elegans

**DOI:** 10.1101/101998

**Authors:** Limin Hao, Yongguang Tong, Kristin Harrington, Jessica L. O’Neill, Afsaneh Sheikholeslami, Xin Wang, Jonathan H. Freedman, Bruce M. Cohen, Edgar A. Buttner

## Abstract

Clozapine binds and has significant effects on multiple neurotransmitter receptors, notably including some dopamine receptors. Downstream of these receptors, clozapine affects the balance of Gi− and Gq-dependent second-messenger signaling. We used *Caenorhabiditis elegans* as a genetic model to study further how clozapine affects both dopamine receptors and downstream Gq mediated signaling. Four of six worm dopamine receptor orthologs, *dop-1, dop-2, dop-4*, and *dop-5* produced resistance to clozapine induced developmental delay when mutated, suggesting that both type I and type II dopamine receptors mediate the behavioral effects of clozapine in *C. elegans*. Beyond these receptors, reduction of function of one of the G proteins, *egl-30* (Gαq), produced greatly increased susceptibility to clozapine. Gαq has multiple known downstream effects. Among these is the control of acetylcholine release, which is in balance with monoamines in the human brain and is another target of clozapine and other antipsychotic drugs. We tested for downstream effects on acetylcholine at the neuromuscular junction upon clozapine treatment but found no evidence for effects of clozapine. In contrast, modulation of Gαq upstream leads to worms that are either more resistant or more susceptible to clozapine, emphasizing the importance of Gαq proteins in mediating effects of clozapine. A genetic screen for suppressors of *egl-30* recovered eight mutants. By characterizing the behavioral effects of these mutants, we found that clozapine exerts its function on development by affecting Gαq signaling through control of the pharyngeal pumping rate. A whole-genome sequencing technique was utilized and identified a list of candidate genes for these suppressor mutations. Further characterization of these mutants promises the discovery of novel components participating in Gαq signaling and a better understanding of the mechanisms of action of clozapine.

## Introduction

Clozapine is an atypical antipsychotic drug (APD), mainly used for schizophrenia that does not improve following the use of standard antipsychotic medications. The pharmacology of clozapine shares some features with other APDs, but other features are unique, and its sites of action are especially complex.

Standard antipsychotic drugs are thought to achieve some of their important effects (both beneficial and unwanted) by acting as antagonists or partial agonists at D2 receptors (JONES AND PILOWSKY 2002). In humans, there are five distinct G-protein coupled dopamine (DA) receptors, and they are divided into two major subtypes: D1-like (D1 and D5) and D2-like (D2, D3 and D4) (CUMMING 2011). These two classifications were originally assigned based on the receptor’s effect on cyclic AMP levels, but also reflect genetic homologies. Selective pharmacological agents and receptor knockouts have been applied to demonstrate that signaling through D1-like and D2-like receptors have opposite effects on many aspects of cell activation and some behaviors (KELLY et al. 1998; GONG et al. 1999; MCNAMARA et al. 2003). However, simultaneous stimulation of both D1 and D2 type receptors shows that these receptors also have synergistic effects on certain behaviors (PLAZNIK et al. 1989; GONG et al. 1999). By comparison to standard antipsychotic medications, clozapine binds weakly to D2, more strongly to D4, among dopamine type II receptors, and most strongly to Type I dopamine receptors, including the D1 receptor. It also binds with high affinity to histamine (H1), acetylcholine muscarinic (M1), α-adrenergic (α1), and multiple serotonin (5-HT2A, 5-HT2C, 5-HT7) receptors (NAHEED AND GREEN 2001).

The *Caenorhabditis elegans* (*C. elegans)* genome encodes six genes for dopamine receptors, and four of these genes have been validated as orthologous to human DA genes, based on sequence similarities, pharmacological profiles, and biochemical properties (SUO et al. 2002; SUO et al. 2003; CHASE et al 2004; SUGIURA et al. 2005). As in humans, these receptors are divided into two categories with opposite effects in modulating cyclic AMP levels; agonists at *dop-1* and *dop-4* upregulate cyclic AMP levels and agonists at *dop-2* and *dop-3* do the opposite. The other two orthologs, T02E9.3 and C24A8.1, named *dop-5* and *dop-6*, have similarities to the *dop-3* sequence, but they have not yet been experimentally validated to be D2-like. Signaling transduction mediated by these dopamine receptors controls multiple worm behaviors, including locomotion, egg laying, defecation, basal motor activity, sensation/response to food sources, and habituation to touch (MCDONALD et al. 2006). DOP-1 and DOP-3 have antagonistic effects on locomotion, by acting in the same motor neurons, which co-express the receptors and which are not postsynaptic to dopaminergic neurons.

Direct effects on DA and other receptors are the start of a cascade of reactions which next passes through G proteins, as second messengers. A non-biased genetic screen identified two antagonistic G proteins, Gαo and Gαq, activated by these receptors and mediating their downstream effects (CHASE et al. 2004). Gα-GTP and Gβγ dimers transmit receptor-generated signals to downstream effector molecules and protein-binding partners until the intrinsic GTPase activity of Gα hydrolyzes GTP to GDP and the inactive subunits re-associate. G proteins are localized and function primarily at the plasma membrane, in contact with cell surface receptors, but also function intracellularly at endosomes (SLESSAREVA et al. 2006)., Gαq activates phospholipase C-β (PLCβ), thereby generating the second messengers inositol-1,4,5-triphosphate

(IP3) and diacylglycerol (DAG). These, in turn, release stored Ca^2+^ and activate protein kinase C. Gαq also activates GEF p63RhoGEF (ARHGEF25) and its homologs Trio and Duet. These tertiary messengers subsequently modulate numerous enzymatic processes, metabolic pathways, components of the cytoskeleton and the release of extracellular signaling molecules.

*C. elegans* has 21 Gα, 2 Gβ and 2 Gγ genes and it expresses one ortholog of each of the four mammalian families GSA-1(Gs), GOA-1(Gi/o), EGL-30 (Gq) and GPA-12 (G12). The remaining *C. elegans* Gα subunits do not share sufficient homology to mammalian Gα proteins to allow classification (BASTIANI AND MENDEL 2006). They may play roles similar to the four types of G proteins, e.g., GPA-14 is able to interact with DOP-2 and shows close similarity to inhibitory G proteins that modulate anterior touch habituation and chemosensory associative conditioning (PANDEY AND HARBINDER 2012; MERSHA et al. 2013). In a fully studied *C. elegans* neural muscular junction, it has been discovered that a network of G protein signaling pathways controls the release of synaptic vesicles and/or dense-core vesicles (PEREZ-MANSILLA AND NURRISH 2009). The Gαq, Gα12 and Gαo pathways converge to control production and destruction of the lipid-bound second messenger DAG at sites of neurotransmitter release. DAG subsequently acts via at least two effectors, MUNC13 and PKC, to control the release of both neurotransmitters and neuropeptides from motor neurons. The Gαs pathway converges with three other heterotrimeric G-protein pathways downstream of DAG to regulate neuropeptide release (BASTIANI AND MENDEL 2006).

G protein mediated effects of clozapine have been studied in cultured cells and mammals. In cultured PC12 cells, clozapine downregulates the expression of tyrosine hydroxylase. This effect can be abolished by treatment with N-ethylmaleimide, a reagent that decouples Gi/o protein from its adjunct receptors, suggesting that clozapine modulates Gi/o protein signaling pathways (TEJEDOR-REAL et al. 2005). In the rat brain, the serotonin receptor, 5-HT2A, recruits Gq proteins promoting PLC stimulation, and clozapine antagonizes 5-HT-induced phosphoinositide (PI) hydrolysis, probably by inhibiting Gq proteins (MANNOURY LA COUR et al. 2009). Recently, it has been found that clozapine binds the mGluR2/2AR receptor heterocomplex, which also alters the balance of Gi− and Gq-dependent signaling (FRIBOURG et al. 2011).

In our laboratory, we have used the well-documented clozapine-induced developmental delay/lethality phenotype in *C. elegans* to study the mechanisms of action of clozapine. We performed a genome-wide RNAi screen and identified 40 genes suppressing the effects of clozapine and characterized some of these genes (SAUR et al. 2013; WANG et al. 2014; HAO et al.2016). To increase understanding of specific mechanisms underlying these findings, we now take advantage of the known effects of clozapine on dopamine receptors and extend them to the study of downstream effects on G protein signaling pathways. The new findings suggest that a known G protein signaling downstream pathway is unaffected, but a non-biased genetic screen revealed new G protein pathway components that could mediate effects of clozapine.

## Materials and Methods

### Nematode growth and worm strains

Nematodes were cultured on NGM plates in a 20°C incubator (LEWIS AND FLEMING 1995). The wild type N2 worm was used in these experiments as a control species. Other worm strains used are listed as follows: DA1077: *egl-30(ad810); dpy-5(e61)/szT1[lon-2(e678)];*+/*szT1*, EAB66-71 and EAB73-74: suppressors of *egl-30(n686)*, LX645: *dop-1(vs100)*, LX702: *dop-2(vs105)*, LX703: *dop-3(vs106)*, LX704: *dop-2(vs105); dop-3(vs106)*, LX705: *dop-1(vs100); dop-3(vs106)*,
*LX706: dop-1(vs100); dop-2(vs105)*,LX734: *dop-1(vs100); dop-2(vs105); dop-3(vs106)*, MT1434: *egl-30(686)*, MT2609: *egl-30(n715n1190)*, NM1380: *egl-30(js126)*, PS2444: *dpy-20(e1282); syIs36*, PS4263: *egl-30(md186); dpy-20(e1282); syIs105*, RB1254: *dop-4(ok1321)*,RB1680: *dop-6(ok2090)*, RB785: *dop-5(ok568)*. The other worm strains employed can be seen in Table 1.

### Developmental drug assay

The developmental drug assay was performed as previously described in (HAO AND BUTTNER 2014). In brief, well-grown adult animals were treated in bleach solution (hypochlorite:1 N NaOH: ddH2O(1:4:5)) to obtain synchronized eggs. 25 synchronized eggs were then placed onto 12-well NGM assay plates containing variable concentrations of drugs. The drugs were initially dissolved in DMSO to obtain 80 mM stock solutions, which were diluted into 1.7 mM acetic acid. Treated worms were transferred onto the NGM plates. Every 24 hours thereafter, the plates were observed and the developmental status of the worms was recorded. For each worm strain, the experiment was repeated at least three times. The concentration of clozapine was adjusted according to the sensitivity of the tested worm strain. We present results from the third day of observation, as results at this time optimally identify and represent the phenotype of strains with different growth characteristics.

### Worm Behavioral assays

The worms used for behavioral assays were staged young adult worms. Unless stated otherwise, these were obtained by picking well-grown L4 worms onto NGM plates 24 h before the assay to allow them grow to young adulthood. For the pharyngeal pumping assay, ten worms were placed on NGM assay plates containing various concentrations of drugs, allowed to adapt for 30 min, then, the number of contractions of the pharyngeal bulb was counted for 20 s for each worm. For the aldicarb assay, the NGM plates were supplemented with 1 mM aldicarb. Thirty worms were placed onto the assay plates, observed every 30 min and scored as moving or paralyzed worms. Paralyzed worms did not move when poked by the worm pick. For the locomotion assay, ten young adult worms were placed on NGM plates seeded with OP50 *E. coli*. After adaptation for 5 min, the number of wiggles of the body was counted for 30 s for each worm. For the egg retention assay, well-fed young adult worms were picked onto NGM plates with food and allowed to grow for 24 h. The assayed worms were picked onto a slide with an agarose pad with 20 mM sodium azide, and observed under a Zeiss Axio2 microscope using the 100x objective lens. The number of eggs in the belly were counted. All assays were repeated three times and two-tail t-tests were used to compare the mutant strain with the control strain.

### Genetic screen for suppressors of *egl-30(n686)*

Synchronized L4 worms were treated with 50 µM ethyl methanesulfonate and shaken for 4 hrs. The treated worms (P0) were collected and washed twice with M9 buffer and transferred onto two NGM plates to grow for 24 hrs. Ten adult worms were transferred onto each of ten fresh plates and allowed to lay eggs for 4 hrs. Then they were transferred onto another ten fresh plates and allowed to lay eggs (F1) overnight. After they grew to adulthood, ten F1 worms were transferred onto 60 NGM plates containing 250 or 350 µM clozapine, and allowed to lay eggs (F2) overnight. After 1-2 days, ten surviving F2 worms were transferred onto each of 60 NGM plates and allowed to grow to adulthood. Then six F2 worms were transferred onto NGM plates containing 300 µM clozapine (*egl-30(n686)* was also picked as a control) and allowed to lay 25 eggs (F3), after which the adult worms were removed. After 2-3 days, if there were more worms surviving in a plate compared to the *egl-30* control, the plate of worms were classified as likely suppressors. The suppressors were recovered and validated by testing them with a standard clozapine developmental delay assay (HAO AND BUTTNER 2014).

### Whole-genome sequencing and data analysis

Worms were grown on 10-20 8.5 cm NGM plates until they were gravid adults, then washed with M9 buffer into a 50 ml tube and centrifuged at 2000xg for 3 min. The supernatant was removed and 30 ml of bleach solution was added. Then the tube was inverted for 7-10 min until all the worms were dissolved, followed by centrifugation at 1500xg for 1 min to collect eggs. The eggs were washed once with M9 buffer, transferred onto 10 5.5 cm unseeded NGM plates, and incubated overnight to allow the eggs to hatch. The hatched L1 worms were washed off the plates with M9 buffer and collected by centrifugation at 1500xg for 1 min, then washed twice with M9 buffer. The clean L1 worms were used immediately or stored at −80°C. DNA wasisolated via a Gentra Puregene Kit (Qiagen). The concentration of the final DNA preparation was determined using the Qubit assay (Invitrogen). 1 µg DNA was used for preparing the sequencing library according to the TruSeq®DNA PCR-free sample preparation kit (Illumina). DNA sequencing was performed using an Illumina HiSeq sequencer.

The sequencing results were analyzed at the high-performance computation facility of the Partners Healthcare System. The reads were mapped to the reference genome (WS220) with Bowtie-0.12.7, and the variant calls were done with Samtools. The variants were checked in igv. We only focused on mutations that occurred in coding regions and splicing sites at which variants could potentially cause disturbance of protein structures.

Strains are available upon request.

## Results

### Dopamine receptors mediate effects of clozapine

There are six dopamine receptor orthologs in the *C. elegans* genome, and mutants in each ortholog are available. We performed developmental assays by growing the mutants and wild type worms in plates treated with 0, 300, 400, and 500 µM clozapine. As shown in Fig. 1A, the lethality rate of wild type worms was high and surviving worms were at the L1 stage. By comparison, the lethality rates of *dop-1, dop-2, dop-2; dop-3, dop-1; dop-3, dop-1: dop-2, dop-1; dop-2; dop-3* mutants were much lower and some of the worms developed to the L2 stage. By comparison, the lethality rate of *dop-3* mutants was as high as that of the wild type strain. These observations, that *dop-1* and *dop-2* mutants were resistant to clozapine, while *dop-3* mutants were not resistant to clozapine, suggest that both *dop-1* and *dop-2* mediate effects of clozapine, but *dop-3* does not. We also tested mutants of the other three dopamine receptors, *dop-4, dop-5* and *dop-6*, and found that *dop-4* and *dop-5* mutants were resistant to the effects of clozapine, but *dop-6* mutants were not (Fig. 1B).

**Fig. 1.**
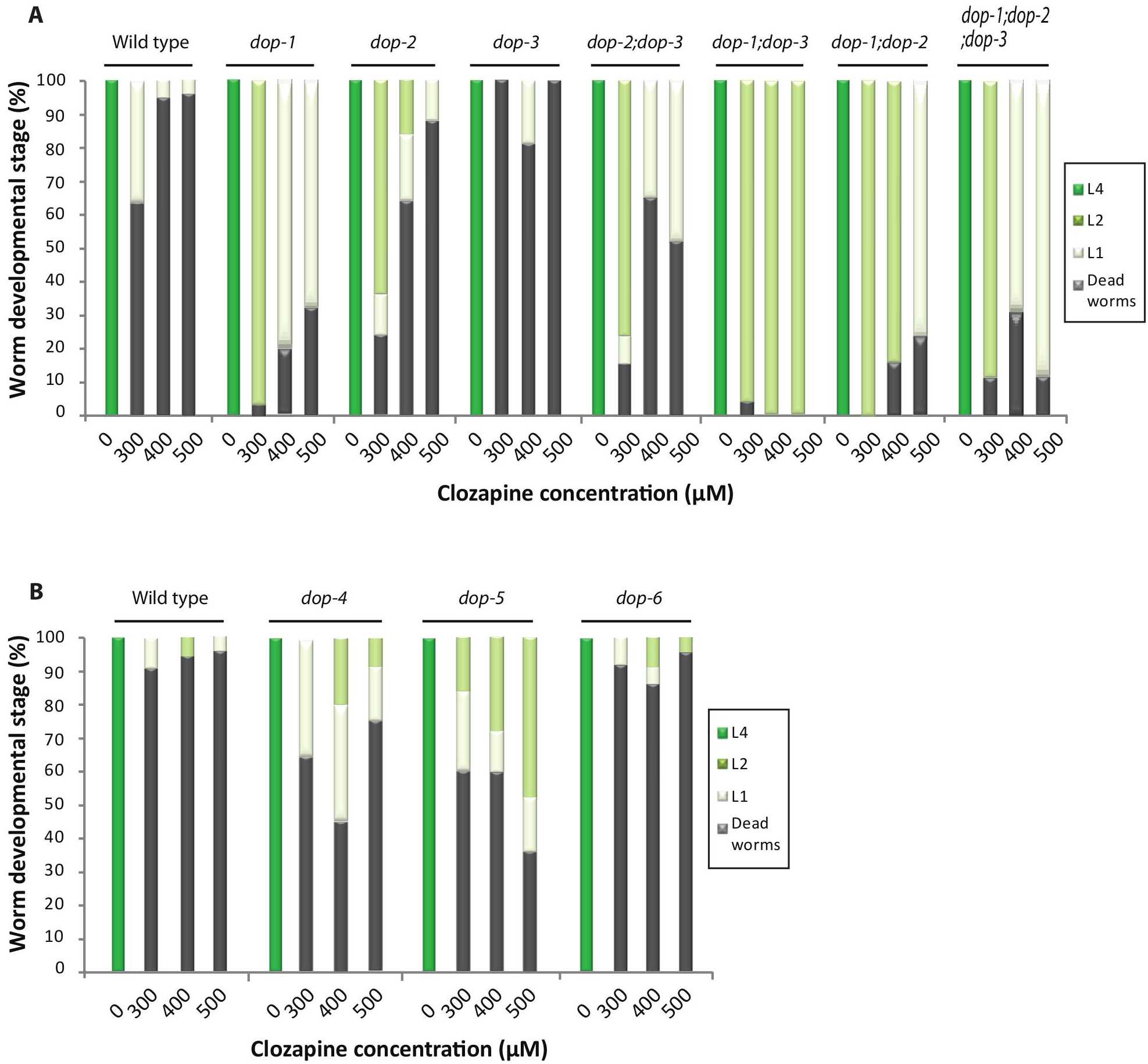
Dopamine receptors mediate clozapine effects. **(A)** Dopamine receptor mutantss,dop-1(vs100), dop-2(vs105), dop-3(vs106), and combinations of these mutants, dop-2(vs105); dop-3(vs106), dop-1(vs100); dop-3(vs106), dop-2(vs105); dop-3(vs106), dop-1(vs100); dop-2(vs105); dop-3(vs106) were treated with clozapine at 0, 300, 400, 500 µM. **(B)** Dopamine receptor mutants, dop-4(ok1321), dop-5(ok568), and dop-6(ok2090) were treated with clozapine at 0, 300, 400, 500 µM.

### Gαq proteins mediate effects of clozapine

Dopamine receptors are G protein coupled, and the alpha subunit of G protein has been found to be associated with the effects of clozapine in mammalian models. To test for this association in *C. elegans*, we studied mutants of the Gαq ortholog, *egl-30*, in worms. Compared to wild type, the lethality rate of a reduction of function (*rf*) mutant, *egl-30(n686)*, was dramatically increased and development was delayed. Conversely, the lethality rate of a constitutively active mutant, *egl-30(js126)*, was slightly reduced and development was promoted (Fig. 2A). To confirm this observation, we conducted drug assays on two other *rf* mutant alleles, *egl-30(ad810)* and *egl-30(n715n1190)*,and found that they were also susceptible to clozapine (Fig. 2B). We also tested two worm strains carrying the integrated transgene of *egl-30* DNA sequence in either wild type background or an *egl-30* mutation background, supposed to have higher Gαq level than wild type, and found that they both reduced lethality rates and promoted development, compared to wild type (Fig. 2B).

**Fig. 2.**
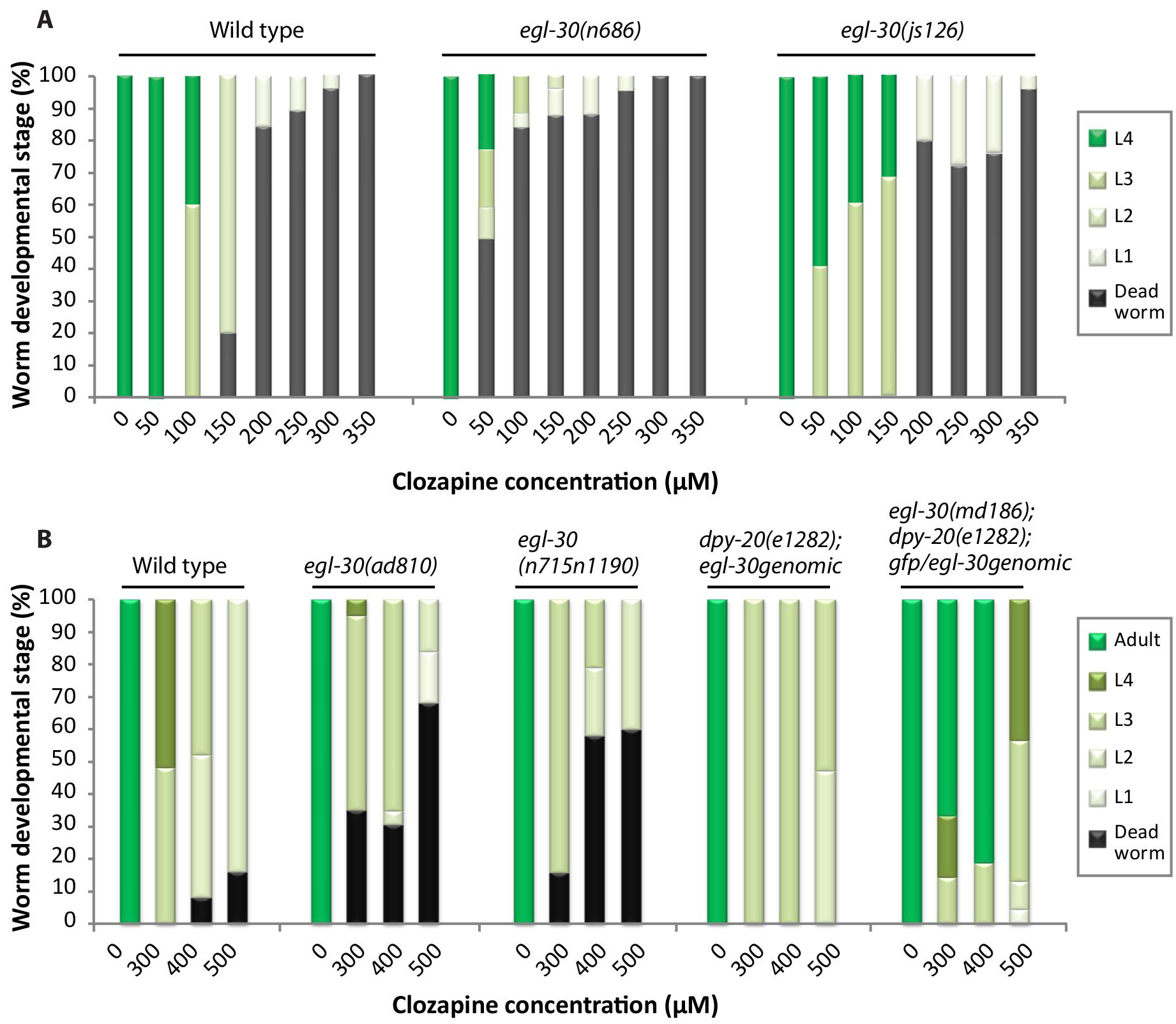
Gαq mediates clozapine effects. (A) Two Gαq mutants, one reduction of function (*rf*)mutant *egl-30(686)* and one constitutive activity mutant *egl-30(js126)*, was compared with that of the wild-type N2 strain by testing them in a series of concentrations of clozapine at 0, 50, 100, 1*50, 200, 250, 300* and *350* µM. (B) Two other *egl-30(rf)* mutants, *ad810, n715n1190*, and two worm strains carrying the over-expressed genomic rescue constructs, *dpy-20(e1282); egl-30* genomic and *egl-30(md186); dpy-20(e1282); gfp/egl-30* genomic were treated with clozapine at 0, 300, 400, 500 µM.

### Various known downstream components of the Gαq signaling transduction pathway are not involved in mediating effects of clozapine

The *egl-30* gene is expressed ubiquitously and required for worm viability, locomotion, egg laying, synaptic transmission and pharyngeal pumping. We used egg laying and locomotion phenotypes to study downstream behavioral effects of changes in Gq signaling pathways. We also studied one pathway related to synaptic transmission. Gα protein-involved signaling controls the release of acetylcholine and other neuropeptides (Fig. 3).

We performed developmental drug assays on three types of Gα proteins, *goa-1* (Gαo), *gpa-12* (Gα12), *gsa-1* (Gαs) and found that their mutants are no different than wild type in response to clozapine (Fig. 3 and Table 1).

**Fig. 3.**
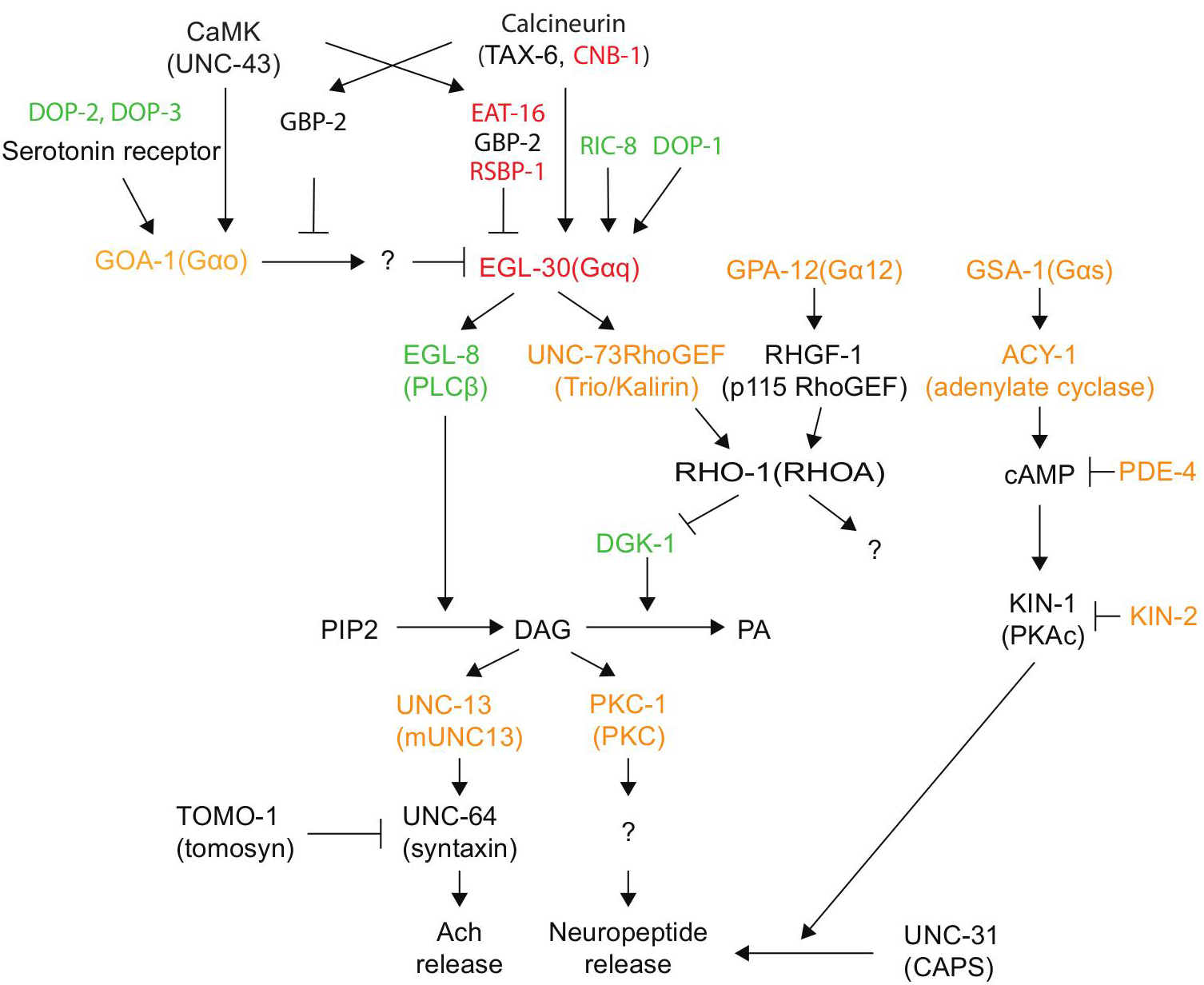
Testing downstream clozapine effects via an established Gα signaling pathway that 452 regulates Acetylcholine receptor and neuropeptide release in *C. elegans*. Testing downstream clozapine effects via an established Gα signaling pathway that regulates Acetylcholine receptor and neuropeptide release in *C. elegans*. Adapted from Perez-Mansilla and Nurrish (2009). The green colored components indicate mutants that are resistant to clozapine; the red colored components indicate mutants that are susceptible to clozapine; and the orange colored components indicate mutants that are phenotypically similar to wild type strains. Mutants in the black components were not tested.

In the Gq and Gαs signaling pathways, we conducted developmental drug assays on the downstream components of the Gq signaling pathway, namely, *egl-8, unc-73, dgk-1, unc-13*,
*pkc-1*, and found that *dgk-1* suppressed the effects of clozapine, while *egl-8* was similar to wild type or slightly suppressed the effects of clozapine. By comparison, *unc-73, unc-13*, and *pkc-1* strains all responded to clozapine the same as wild type strains (Table 1). In addition, developmental drug assays performed on the downstream components of the Gαs signaling pathway, *acy-1, pde-4*, and *kin-2*, showed that they all responded to clozapine the same as wild type strains (Table 1). These observations suggested that these known downstream components of Gq and other Gα signaling pathways were not involved in mediating the effects of clozapine.

### Gαq modulators alter effects of clozapine

We investigated the role of modulators of Gαq by conducting developmental assays and discovered that mutants of *ric-8*, a non-receptor activator of Gαq in both *C. elegans* and mammals, are resistant to clozapine (Fig. 3, Table 1). Mutants of three other modulators of Gαq, *cnb-1, eat-16*, and *rsbp-1* were found to be more susceptible to clozapine (Table 1).

### Isolation and characterization of Gαq mutant suppressors of clozapine effects

We performed a forward genetic screen for suppressors of *egl-30(n686)*, taking advantage of the hypersensitivity of *egl-30(n686)* to clozapine. We found eight mutants that were more resistant to clozapine than *egl-30(n686)*, and these suppressors were named EAB66-71 and EAB73-74. The strongest suppressor was EAB66, which was even more resistant to clozapine than wild type (data not shown). We characterized these mutants for other behavioral phenotypes and compared them to wild type and *egl-30(n686)* worms. Aldicarb is an inhibitor of cholinesterase, causing rapid accumulation of acetylcholine at the synaptic cleft, and thereby paralysis of the worms. The *egl-30(n686)* mutant is resistant to aldicarb compared to wild type. The suppressor EAB66 is phenotypically like the wild type worm, whereas EAB67 remains phenotypically like the *egl-30(n686)* worm (Fig. 4A). In the locomotion assay, strains carrying the suppressor EAB66 mov faster than *egl-30(n686)*, but they do not move at the wild type level; whereas the EAB67 strain is even slower than *egl-30(n686)* (Fig. 4B). As for the egg retention assay, EAB66 and EAB67 strains are both more similar to *egl-30(n686)* strains than wild type worms (Fig. 4C). However, in the pharyngeal pumping assay, all the suppressors show activity that is statistically significantly different from *egl-30(n686)*, except EAB73 at 450 µM (Fig. 4D).

**Fig. 4.**
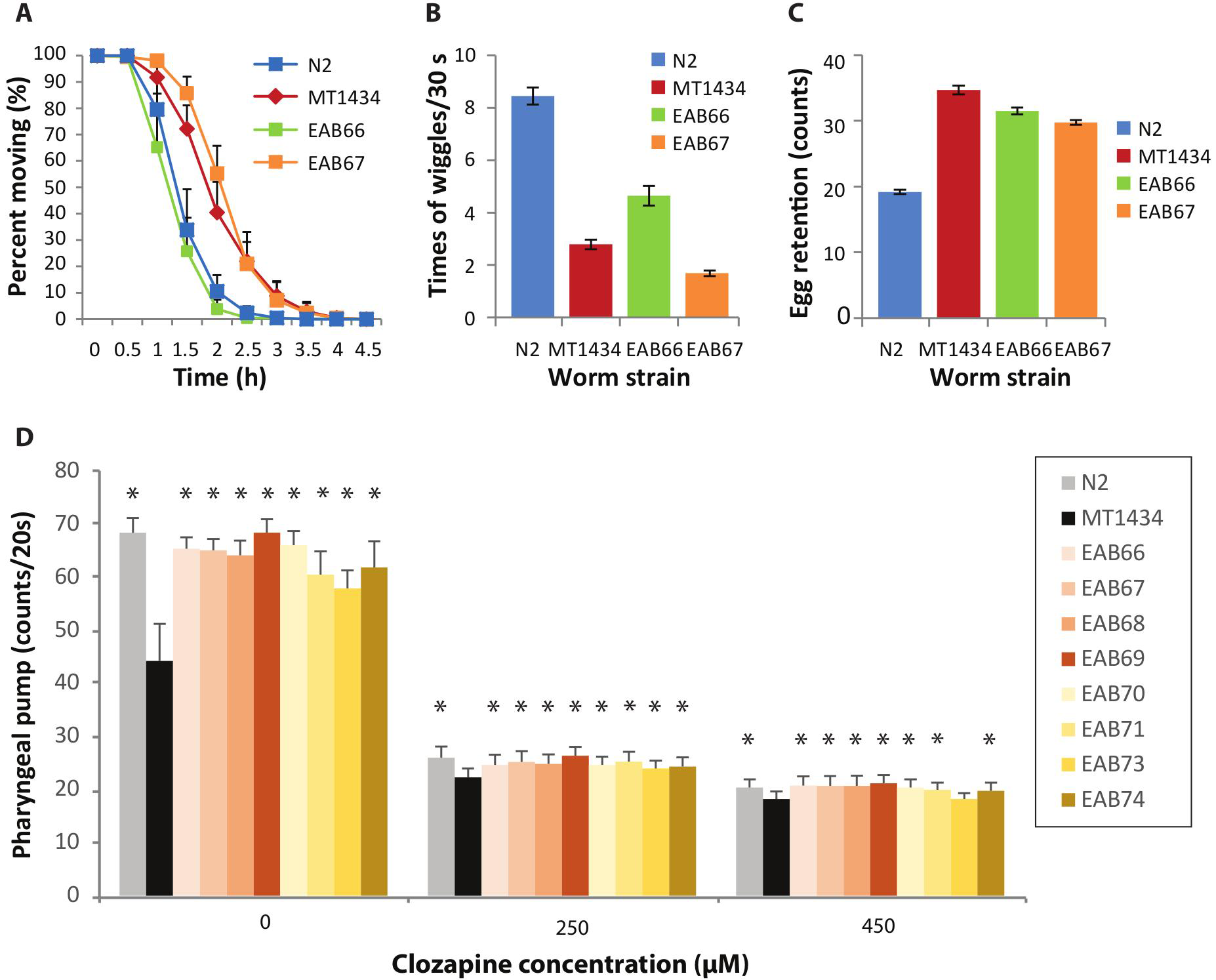
Characterization of the *egl-30* suppressors. Characterization of the egl-30 suppressors. (A) N2, MT1434 and two EAB66-67suppressor strains were treated with 1 µM aldicarb and examined for movement at 0, 0.5, 1, 1.5, 2, 2.5, 3, 3.5, 4, 4.5 hrs. The data are from triplicated experiments with total numbers of tested worms N=70. (B) Locomotion assays of the N2, MT1434 and EAB66-67 suppressor strains. There are statistically significant differences in phenotype among these four strains, P<0.001. The data are from triplicated experiments with a total number of tested worms N=35. (C) Egg retention assays of the wild type N2, MT1434 and EAB66-67 strains. Comparison of these four strains again shows statistically significant differences, P<0.05. The data are from triplicated experiments with a total number of tested worms N=60. (D) Pharyngeal pumping assays of wild type (grey bar), MT1434 (black bar) and eight MT1434 suppressor strains (EAB66-71, 73, 74) (color bars). Comparisons between each of the suppressors and the MT1434 strain show statistically significant differences, but differences are not observed between the suppressor bearing strains and wild type strains. (* indicates p<0.001 between each of the mutants and MT1434.) The data are from triplicated experiments with a total number of tested worms N=30.

**Table 1.** Summary of drug assays on worm strains in 473 known C. elegans Gα signaling transduction. +:Weak suppressor, ++: Intermediate suppressor, +++: Strong suppressor, −:Weak enhancer, −−: Intermediate enhancer, −−−: Strong enhancer, ±: Wild-type.

| Ortholog in human | Gene | Allele | Strain | Developmental assays |  |  |  | Concluded phenotype |
| --- | --- | --- | --- | --- | --- | --- | --- | --- |
| Gα<sub>0</sub> | *goa-1* | *sa734* | DG1856 | ± | ± | -- | | WT |
| PLC β | *egl-8* | *n488* | MT1083 | ++ | + | ± | | Suppressor |
| HMUNC13 | *unc-13* | *e51* | MT7929 | + | -- | ± | | WT |
| DGKQ | *dgk-1* | *sy428* | PS2627 | ++ | +++ | + |  | Suppressor |
| PRKCE | *pkc-1* | *ok563* | RB781 | -- | -- | ++ | + | WT |
| GNEF | *unc-73* | *e936* | CB936 | + | ± | ± | | WT |
| RGS6,7,9,11 | *eat-16* | *sa609* | JT609 | ± | --- | --- | - | Enhancer |
| Calcineurin B | *cnb-1* | *jh103* | KJ300 | -- | -- | -- | -- | Enhancer |
| None | *rsbp-1* | *vs163* | LX1270 | --- | -- | --- |  | Enhancer |
| Synembryn-A | *ric-8* | *md303* | RM1702 | ++ | +++ | ++ |  | Suppressor |
| GNA12 and 13 | *gpa-12* | *pk322* | NL594 | ± | ± | ± | | WT |
| GNAL and S | *gsa-1* | *ce81* | KG421 | ± | ± | ± | | WT |
| ADCY9 | *acy-1* | *ce2* | KG518 | ± | ± | ± | | WT |
| PDE4A-D | *pde-4* | *ce268* | KG744 | ± | ± | ± | | WT |
| PRKAR1A-B, CNBD1 | *kin-2* | *ce179* | KG532 | ± | ± | ± | | WT |

**Table 2.** Candidate mutated genes of egl-30(n686) 477 suppressors revealed by whole-genome sequencing.

| Suppressor | Location | Gene name or putative function | Site | nn change | AA change |
| --- | --- | --- | --- | --- | --- |
| EAB66 | F08G5.7 /IV | *clec-184* | 12433358 | C/T | Q/Stop |
|  | M02b1.1/IV | *srf-3* | 12850684 | C/T | Q/Stop |
|  | C31B8.7/V | No info, only in nematodes | 2901829 | C/T | Splicing site |
|  | T08A9.1/X | *atg-11* | 7339350 | C/T | Q/Stop |
| EAB67 | T13B5.7/II | *lips-13* | 1115919 | G/A | Q/Stop |
|  | C41A3.1/V | *fasn-1* (fatty acid synthase) | 5430972 | C/T | Q/Stop |
| EAB68 | Y39H10A.4/V | Not known function; Only in nematode | 3733499 | C/T | Splicing site |
|  | *srj-27/V* | Predicted to have transmembrane signaling receptor activity, based on protein domain information. Only exists in nematode | 15126479 | G/A | Q/Stop |
|  | zc13.10/X | Not known function; Only in nematode | 902597 | T/C | Initial M/V |
|  | F21G4.3/X | Not known function; Only in nematode | 9896512 | C/T | R/Stop |
|  | zc506.1/X | Ortholog of human EDEM3 (ER degradation enhancer); predicted to have mannosyl-oligosaccharide 1,2-alpha-mannosidase activity and calcium ion binding activity, based on protein domain information. | 9963398 | C/T | Q/Stop |
| EAB69 | *clec-131/I* | C-type lectin, only existed in nematode. | 3575866 | C/T | W/Stop |
|  | *rmd-3/I* | Ortholog of human RMDN2 and RMDN3 (regulator of microtubule dynamics). | 11339397 | C/T | Splicing site |
| EAB70 | Y87G2A.7/I | *nyn-2*, involved in RNAi | 13574346 | C/T | Q/Stop |
|  | Y66D12A.14 /III | Amino adipate-semialdehyde dehydrogenase | 11544129 | C/T | Q/Stop |
| EAB71 | ZK1055.7/V | Involved in innate immune response | 6603050 | G/A | Q/Stop |
|  | C41A3.1/X | Ortholog of human FASN (fatty acid synthase). | 5430972 | C/T | Q/Stop |
| EAB73 | *unc-89* | Encodes several protein isoforms variably characterized by the presence or absence of SH3, DH, PH, immunoglobulin and myosin light chain kinase (MLCK)-like protein kinase domains | 4082080 | A/T | R/Stop |
|  | Y54B9A.1 | Kinesin-like protein | 8078419 | G/A | W/Stop |
| EAB74 | *srv-17* | 7TM GPCR, serpentine receptor class v (Srv) | 5869831 | C/T | Splicing site |
|  | F23D12.2 | Encodes a protein containing an F-box and FTH/DUF38 motifs, mediating protein-protein interaction. | 14416566 | G/A | R/Stop |

### Candidate genes for Gαq suppressors revealed by whole-genome sequencing

To identify the specific genes coding for the suppressor mutants, we performed whole-genome sequencing of these mutants and compared the sequences of wild type and *egl-30(n686)* worms. As shown in Table 2, there are two to five mutations occurring in these mutants, each of which changes protein coding sequences. Most of the mutations introduce a stop codon in a coding region; but there are four mutations that change splicing sites; and one mutation changes the initial start codon (M/V) in EAB68 (Table 2).

## Discussion

In the worm model, dopamine receptors participate in mediating the effects of clozapine in producing developmental delay/lethality. Interestingly, when mutated, four out of six dopamine receptors, including both type I and type II dopamine receptors, relieve clozapine-induced developmental phenotypes (Fig. 1). These findings cannot be explained by the simple model in which the two types of dopamine receptors have purely opposite and competing effects at the level of production of cAMP, through activation of Gαs and Gαo. Our observation that neither Gαs nor Gαo mutants display behavioral differences from wild type strains in drug assays is also consistent with a more complex interaction of dopamine receptors and G proteins (Table 1). Notably, there is growing evidence that G protein coupled receptors (GPCRs) execute their functions through oligomerization, including homomerization and heteromerization (FERRE AND FRANCO 2010). Oligomerization allows receptors to achieve new properties, including changes in ligand recognition, G protein-coupling and trafficking (FERRE AND FRANCO 2010. For example, while the D1-D3 receptor heteromer couples to Gs (FIORENTINI et al. 2010), the D1-D2 and D2-D5 heteromers couple preferentially to Gq proteins (HASBI et al. 2010). Clozapine may exert its effects on development through interactions with and modulation of such oligomers of dopamine receptors.

Based on our genetic studies on the components in the four G protein-involved signaling networks (Fig. 2, Table 1), only Gq and its modulators appear to strongly mediate the effects of clozapine. The classical model that Gs and Go balance cAMP production and metabolism is not supported by our findings, since the mutants of Gs and its downstream components do not differ from wild type in their drug response. Gq and Go may form a signaling balance as suggested by Fribourg et al. (FRIBOURG et al. 2011). A Go/Gq heterotrimeric G protein signaling network expressed throughout the nervous system has been shown to regulate locomotion rate, partly by affecting serotonergic neurotransmission. And loss-of-function (*lf*) mutations in *goa-1* (Gαo) cause hyperactivity, whereas *lf* mutations in *egl-30* (Gαq) cause severe lethargy (MILLER et al. 1999; NURRISH et al. 1999). However, while *egl-30* Gαq mutants are highly susceptible to clozapine (Fig. 2), the *goa-1* (Gαo) mutant does not display a resistant phenotype upon clozapine treatment (Table 1). Furthermore, the downstream components, i.e., in the PLC signaling pathway that facilitates synaptic transmission by body-wall muscle motor neurons, do not appear to mediate clozapine effects, as their mutants have the same response as wild type strains (Table 1). Taken together, these observations indicate that the effects of clozapine on worm development are not mediated by these known Gαq downstream signaling elements.

To explore how Gq proteins mediate the effects of clozapine, we performed a genetic screen and identified eight suppressing mutants that were more resistant to the behavioral effects of clozapine than the *rf* of *egl-30* strain. Since the *rf* of *egl-30* exhibited phenotypes including decreased pharyngeal pumping and locomotion rates, reduced sensitivity to aldicarb, and high retention of eggs, we tested whether these behaviors were affected by the suppressing mutations. We tested two suppressors, EAB66 and EAB67, and found that aldicarb sensitivity, locomotion rate and egg retention were changed at baseline in ways similar to *egl*-30, but EAB66 and EAB67 did not have the characteristics of *egl*-30 in altering response to clozapine (Fig. 4). The genes affected by these suppressors might not be involved in the developmental/lethality phenotypes induced by clozapine. In contrast, the pharyngeal pumping rates of all the suppressors recovered to the wild type levels (Fig. 4), indicating that this behavior does correspond to the effects of clozapine on development, which is in agreement with our previously reported findings (HAO et al. 2016).

Whole-genome sequencing revealed that each mutant strain carried 2-5 mutations that would likely change protein structures. Most of the candidate genes in these suppressors are novel. Further studies may identify the components that mediate the effects of clozapine through the Gq protein.

In summary, we studied the actions of clozapine, using *C. elegans* as a genetic model, and observed that both type I and type II dopamine receptors mediate the behavioral effects of clozapine. Further, the results suggest that dopamine receptors form oligomers to signal through Gq proteins. We also found that clozapine exerts its function on development by modulating Gq signaling, which controls the pharyngeal pumping rate. The mutants we identified can be studied further and those studies promise more interesting discoveries involving Gq signaling and a better understanding of the mechanism of action and effects of clozapine beyond its direct effects on cell surface receptors.

## Acknowledgements

Some strains were provided by the CGC, which is funded by the NIH Office of Research Infrastructure Programs (P40 OD010440). We thank Dr. Michael Koelle at Yale University, Dr. Masahiro Tomioka at University of Tokyo, Dr. Paul W. Sternberg at California Institute of Technology, Satoshi Suo at University of Tokyo at Komaba for providing worm strains and other materials. We thank the Enterprise Research Infrastructure 	 Services (ERIS) at Partners HealthCare for their support in the genome analysis. This work is dedicated to the memory of Dr. Edgar (Ned) A. Buttner, who passed away on October 15, 2015, for his inspiring enthusiasm, warm collegiality and excellence in science.

The work was supported by an NIH Clinical Scientist Development award, K08NS002083, funds of the Program for Neuropsychiatric Research, at McLean Hospital, and a NARSAD Young Investigator Award to Edgar A. Buttner.

